# Fallopian Tube Organoid-Integrated Microfluidic Platform for Real-Time Imaging and Outflow Analysis

**DOI:** 10.64898/2026.10.24.761554

**Authors:** Dolan Evans, Heidi Savey-Bennett, Kimberley Perkins, Alice Thompson, Daniel J. Netherwood, Eleanor Doman, Christine K. Schmidt, Peter T. Ruane, Raveen Tank

## Abstract

The fallopian tube is a dynamic, hormonally regulated epithelial conduit whose luminal microenvironment governs oocyte transport, fertilisation, early embryo development, and the initiation of gynaecological diseases including ectopic pregnancy, hydrosalpinx, and serous tubal intraepithelial carcinoma (STIC), yet existing models such as 2D monolayers, intact organoids, and static Transwell systems fail to recapitulate a polarised epithelium lining a perfused lumen under physiological fluid shear while preserving native organoid secretory function. We present a robust and accessible workflow integrating mechanically opened fallopian tube organoids into a custom-designed polydimethylsiloxane (PDMS) microfluidic device, in which organoid fragments form epithelial sheets and re-establish luminal-like architecture under controlled perfusion. The focus of this study is organoid integration and the characterisation of the resulting organoid-derived epithelium, rather than establishing chip cultures from individual cell lines. As this epithelium is built from expandable organoid stock rather than primary tissue, which typically survives only a few days in culture, the approach removes a major bottleneck to establishing organ-on-chip models, and we maintain viable cultures for up to 10 days. Epithelial sheets reproducibly covered more than 90% of the channel area by day 6 (n = 5 devices) and formed continuous ZO-1-positive tight junctions. The platform supports longitudinal brightfield and fluorescence microscopy at subcellular resolution, continuous collection of conditioned flow-through for ELISA, and quantitative analysis of cell coverage and health using CellProfiler software. We characterise the organoid-derived epithelium by benchmarking against tissue culture (TC) formats, comparing nuclear morphology on-chip with conventional well culture, and showing that the damage-associated cytokine interleukin-1α (IL-1α) was detectable in organoid lysates but below the assay limit of detection in both chip outflow and intact-organoid secretions. These findings indicate that culturing mechanically opened organoids on-chip did not produce a detectable increase in IL-1α release relative to intact organoids. Computational flow modelling characterises perfusion through the device and predicts how factors secreted by healthy epithelium and by tumour cells are distributed along the channel. The feasibility of tumour– epithelium co-culture was explored using OVCAR3 ovarian cancer spheroids as a proof-of-concept model to investigate interactions between tumour cells and the fallopian tube epithelium. Together, these results establish the technical feasibility of the platform and support its further development as an accessible system for studying reproductive health conditions affecting the fallopian tube, including tubal damage from infection, endometriosis, STIC lesions, and occlusion.

## Introduction

The fallopian tube is a hormonally regulated epithelial conduit that supports oocyte pick-up, sperm transport, fertilisation and early embryo development, and its dysfunction underlies a broad range of gynaecological disease. Tubal epithelial damage caused by infection or endometriosis can lead to hydrosalpinx and tubal factor infertility: hydrosalpinx is present in around 10-13% of infertile women when diagnosed by ultrasound and is associated with a 74% relative increase in the risk of pregnancy loss following IVF [2], while endometriosis significantly increases the risk of fallopian tube occlusion compared with women without the disease (35.3% versus 20.4% of patients with at least one occluded tube in a recent cohort) [3]. Ectopic implantation, over 95% of which occurs within the fallopian tube, remains the leading cause of first-trimester maternal death and affects an estimated 1-2% of pregnancies [4]. The tubal secretory epithelium is also now recognised as the principal site of origin of high-grade serous ovarian carcinoma (HGSOC), the most lethal gynaecological malignancy, arising via a precursor lesion termed serous tubal intraepithelial carcinoma (STIC) [5]. Understanding how this epithelium responds to injury, infection and early neoplastic change is therefore directly relevant to fertility preservation, reproductive health and cancer prevention, yet the experimental tools available to study it under physiologically realistic conditions remain limited.

Complex *in vitro* biological models which capture the features of their *in vivo* counterparts have long been sought after in the biomedical sciences. Such models are required to better understand disease mechanisms and are important to drug discovery. Although 2-dimensional mammalian cell models have been hugely successful in aiding our understanding of human health and disease, these systems are limited by the simplicity of their structure. 2D cell culture bears little resemblance to the physiological conditions the models attempt to recapitulate [6]. 3D mammalian cell culture has been shown to fundamentally influence the behaviour of cells [6, 7]. Cell migration, gene expression, cell survival and drug responses are some of the factors influenced by the space in which these cells are cultured [8–12].

Human organoid models have been developed for several organs, including a fallopian tube model [7, 13–15]. Organoids are 3D cell structures which capture more structural complexity than traditional mammalian cell culture, maintaining the differentiation pathways of cells contained in those tissues [14, 16, 17]. To encourage the formation of 3D organoids a 3D hydrogel dome acts as a scaffold [18].

Unfortunately, this dome provides an isotropic environment which lacks controlled fluid dynamics, gradient formation and fine-tuned spatial control, similar to traditional cell culture systems. However, when in a monolayer form this fallopian tube organoid model has been shown to maintain a structure comparable to that of the fallopian tube tissue containing both secretory and ciliated cell types with clear apicobasal polarisation [14]. Integrating such complex organoid monolayers, which maintain the complexity of the tissue of study, with microfluidic platforms, addresses many limitations associated with 2D cell culture of a single cell type and 3D culture of organoids.

Organ-on-chip platforms offer a number of key advantages over more traditional mammalian cell culture techniques, most notably fine control over fluid dynamics, mechanical shear, nutrient and gradient delivery, and the ability to image tissue continuously under physiologically relevant conditions [19]. However, integrating complex 3D organoid structures, rather than dissociated single cells, into microscale channels remains technically challenging, and most existing microfluidic epithelial models rely on cell lines or dissociated primary cells rather than differentiated organoid-derived epithelium [9, 19]. Within the female reproductive tract specifically, microfluidic platforms have so far focused mainly on the endometrium and oviduct: compartmentalised endometrium-on-chip devices have modelled stromal-endothelial interactions and cyclical hormone exposure across an idealised 28-day menstrual cycle [20], while an oviduct-on-chip supporting zygote genome reprogramming used epithelial cell lines under perfusion [9]. To our knowledge, none has yet combined patient-derived fallopian tube organoids with a perfused, imageable chip format.

Here, we present a streamlined method for integrating mechanically opened mature fallopian tube epithelial organoids into PDMS microfluidic devices, where organoid fragments seeded in Matrigel diluted 1:5 in PBS establish planar epithelial sheets. This study focuses on organoid integration and characterisation of the resulting epithelium, rather than on establishing epithelial cultures from primary or immortalised cell lines. This approach achieves uniform epithelial coverage within a perfused channel and enables real-time, high-resolution imaging combined with secretome collection. Because the epithelium is built from expandable organoid stock rather than primary tissue, which typically survives only a few days in culture, this removes a major bottleneck to establishing organ-on-chip models of the fallopian tube. We demonstrate maintenance of organoid-derived epithelial sheets for up to 10 days, supporting longitudinal experiments without the need for fresh patient tissue at every timepoint. The platform uses manual pipette tip-based perfusion, making it accessible with minimal specialist infrastructure. We complement the experimental platform with depth-averaged modelling of flow through the device and COMSOL Multiphysics simulations of flow and secreted-factor transport within the culture channel. The latter predict the relative spatial distribution of factors released by healthy epithelium and discrete tumour sites, providing a framework for interpreting downstream epithelial exposure and transport towards the outlet for effluent sampling. To demonstrate relevance to gynaecological disease, we used co-culture with OVCAR3 ovarian cancer spheroids as a proof of concept for studying early tumour–epithelium interactions with the organoid-derived tubal epithelium. This example illustrates the platform’s potential for broader applications in studying tubal damage, infection, endometriosis, occlusion and early neoplastic change.

## Results

### Characterisation of flow within the device

To understand how medium is distributed across the culture chamber, we modelled flow through the device geometry (Figure 2). Because the device is shallow relative to its width and length, flow was computed using a depth-averaged thin-gap approximation. The rows of closely spaced pillars acted as semi-permeable boundaries, with relatively little flow passing through them, and flow was predominantly directed along the channel length. Nonetheless, within one device width of the inlets, fluid had spread from the three localised inlet regions across the full width of each channel, indicating that cells seeded across the chamber are perfused rather than confined to streams leading directly from the inlets.

### Uniform epithelial sheet coverage in a microfluidic chip

We established a fallopian tube epithelial cell microfluidic chip by mechanically breaking fallopian tube organoids and seeding using the protocol outlined in Figure 1. The aim was to achieve significant coverage of fallopian tube epithelial cell sheets in the device and to ascertain how long the sheets could be maintained in the device. Time-lapse microscopy was done to assess the extent of coverage and track fallopian cell sheet formation and maintenance over a period of 10 days post seeding (Figure 3). Tracking the fallopian tube epithelial cell sheets 24 h post inoculation shows areas of tightly packed epithelial cells which maintain attachment to the device over a period of 10 days (Figure 3).

**Fig. 1:**
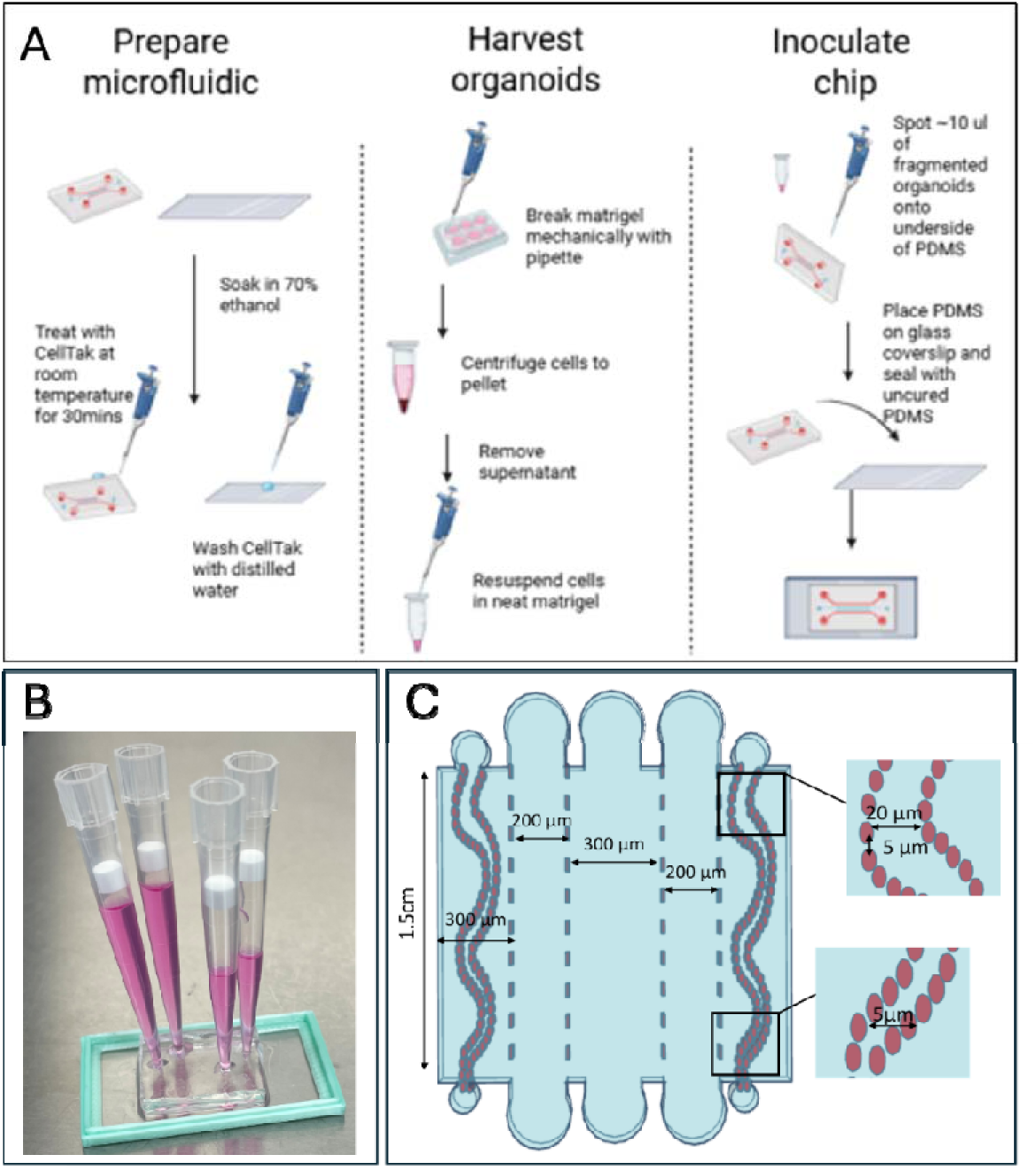
Inoculation of device with fallopian tube organoids using open device inoculation. (A) Overview of the three-stage protocol for seeding fallopian tube organoids onto-chip. In the “Prepare microfluidic” stage, the PDMS device and glass coverslip are soaked in 70% ethanol, treated with CellTak at room temperature for 30 minutes to promote cell adhesion, then washed with distilled water. In the “Harvest organoids” stage, organoids are mechanically fragmented from Matrigel domes using a pipette, pelleted by centrifugation, and the supernatant is removed before resuspending the organoid fragments in Matrigel diluted 1:5 in PBS. In the “Inoculate chip” stage, ∼10 µl of the fragmented organoid suspension is spotted onto the underside of the PDMS device, which is then placed onto the CellTak-coated glass coverslip and sealed with silicone gel around the boundary. (B) Photograph of pipette tips loaded with cell culture medium as reservoirs, with a higher volume at the inlet than the outlet to drive slow passive flow. (C) Schematic of the assembled chip design, showing the central culture chamber (dashed lines) flanked by two curved microfluidic channels (red) for medium perfusion.

**Fig. 2:**
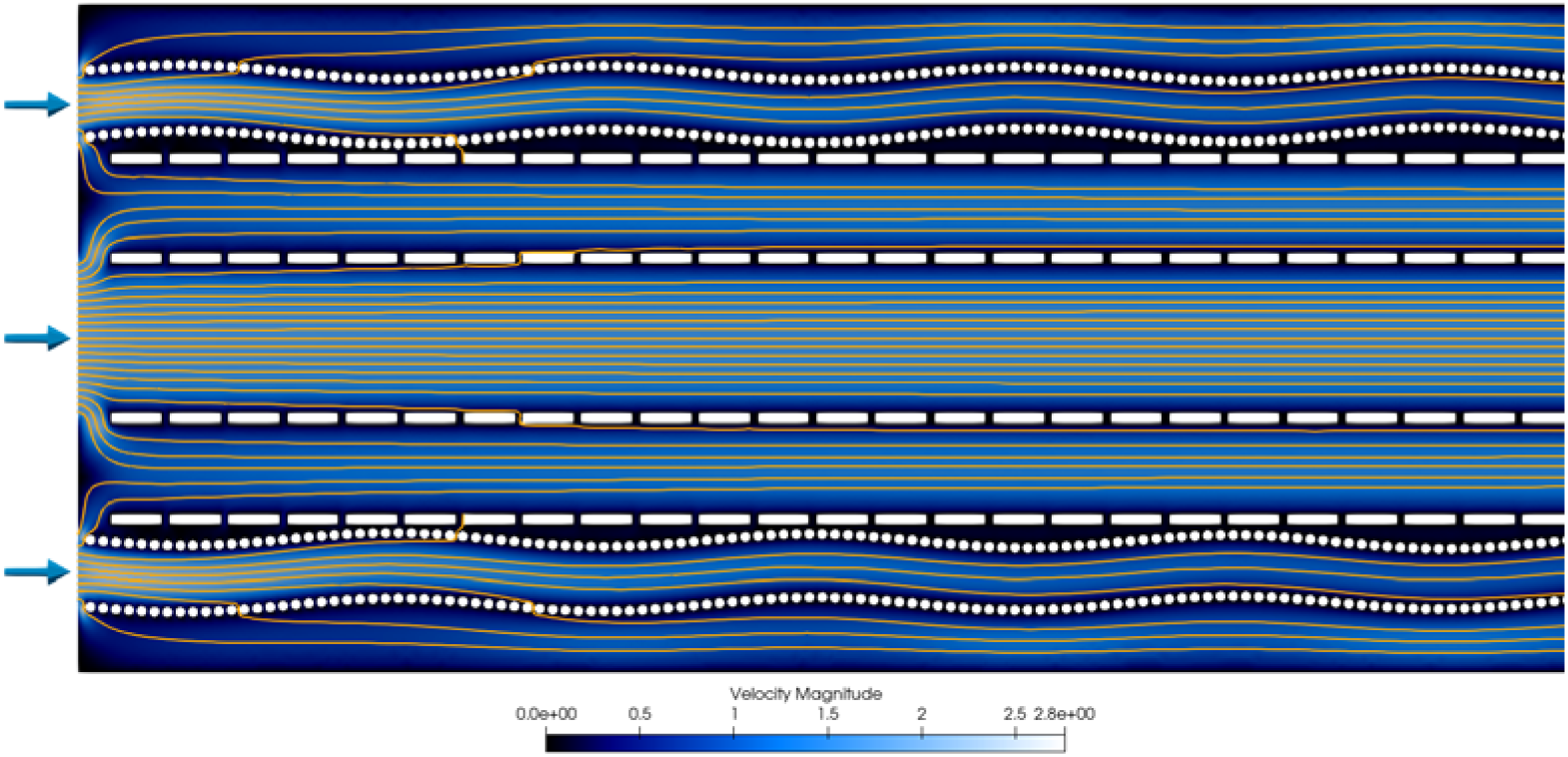
Computational flow model of the device. Two-dimensional streamlines near the start of the channel (top view), computed using a depth-averaged thin-gap model for the device geometry shown. Fluid enters through three inlet regions (arrows). Yellow lines are streamlines, each carrying an equal flux of fluid. The colour map shows relative velocity magnitude, which is zero on all side walls and around the rectangular and circular pillars.

**Fig. 3.**
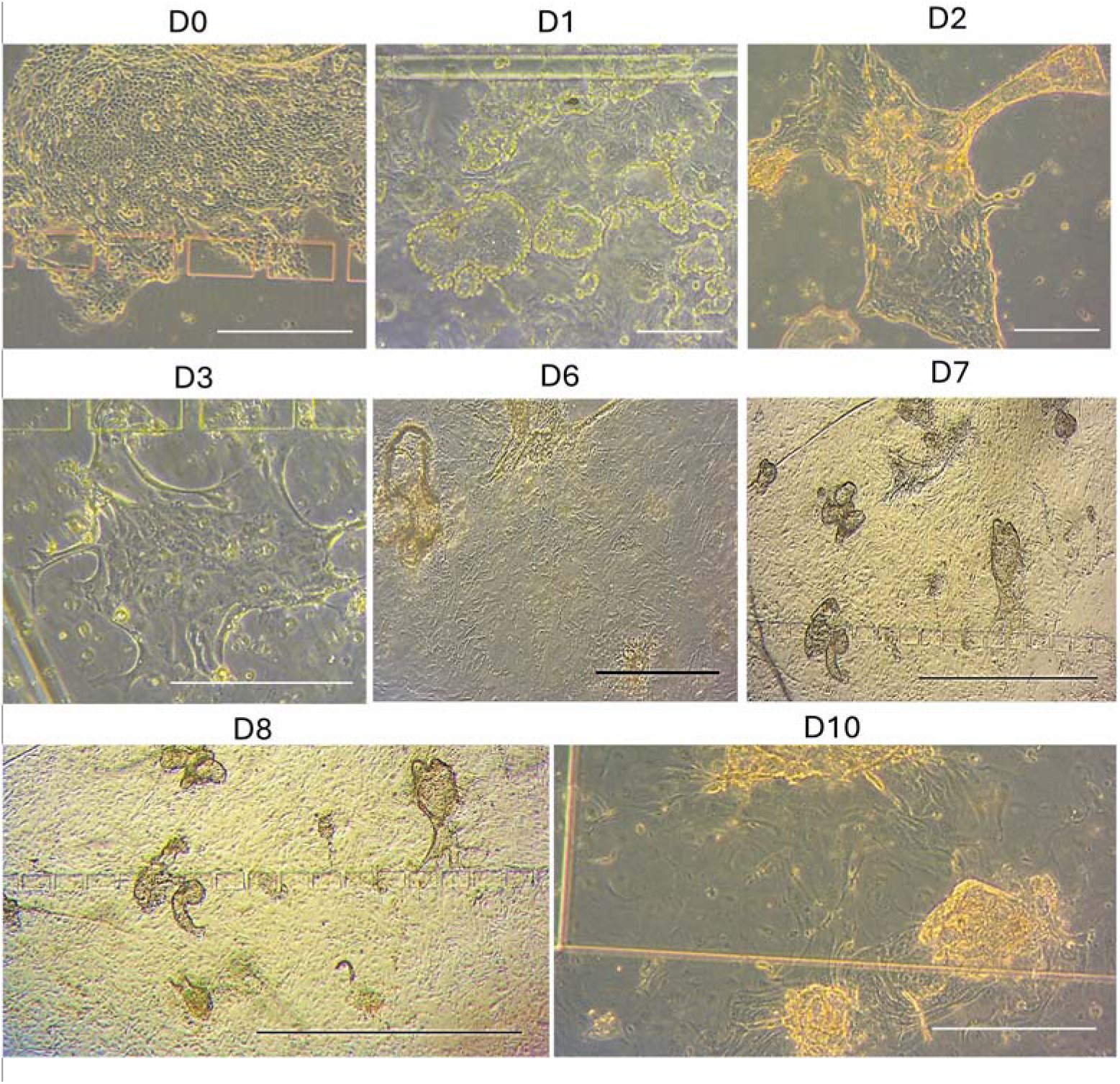
Fallopian tube organoid sheets in a microfluidic device. Time-lapse imaging of fallopian tube epithelial sheet formation in a microfluidic device over a 10-day period post inoculation. Images show one microfluidic device with D0 the day of inoculation and subsequent days. D0-D7 Scale bar 100 µm, D9-D10 scale bar 500 µm. Sheet coverage exceeded 90% of the channel area by day 6 in all devices (n = 5).

Although cell attachment is suggestive of living cells, we wanted to confirm this using a mitochondrial tracker cell viability stain (Figure 4, Bi). Fluorescent green dots only appear in live cells and thus show that the fallopian tube epithelial cell sheets are composed of living cells. Further fluorescent microscopy on fallopian tube epithelial cell sheets in a device 7 days post inoculation shows cells in close proximity with normal nuclear morphology and actin networks highlighting cell-cell interactions (Figure 4Bii). Immunostaining for the tight junction protein ZO-1 showed continuous junctional staining at cell-cell borders across the sheet, indicating that the cells form the tight junctions characteristic of an intact epithelium (Figure 4Ai). As well as cells interacting with one another, cells were also shown to interact spatially with the microfluidic chip. Cells were also shown to make contact through gaps in the chip pillars (Figure 4Aii), demonstrating that chip geometry may influence cell communication and interaction of spatially separated cell populations.

**Fig. 4.**
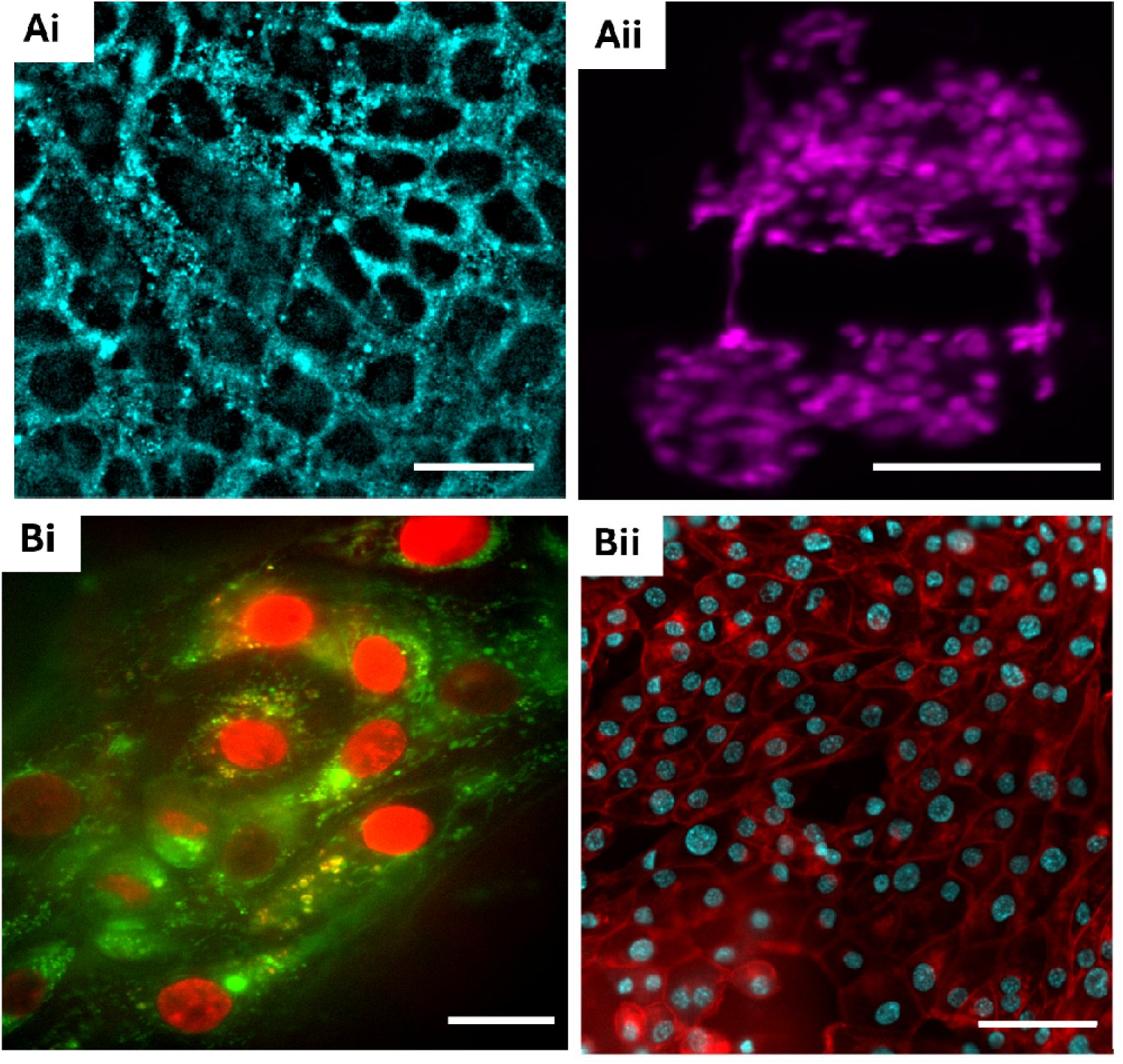
Viability, epithelial organisation and cell–cell interactions in fallopian tube organoid-derived sheets on-chip. Ai) Immunofluorescence image of ZO-1 (tight junctions) in a fallopian tube epithelial sheet on-chip at day 5, showing continuous junctional staining at cell-cell borders. Image acquired using a 60X objective. Scale bar: 20 µm. (Aii) Fluorescence image showing a cluster of cells growing through gaps between the pillars, demonstrating how device geometry can enable communication and interaction between spatially separated cell populations. Scale bar: 50 µm. (Bi) High-resolution image acquired with a 100x oil-immersion objective, showing detailed subcellular organisation and cellular morphology, green is a mito tracker stain for cell viability. Scale bar: 10 µm. (Bii) Fluorescence image acquired using a 20x air objective showing a monolayer grown from inoculated organoid sheets after 1 week in culture. Actin is stained red, highlighting tight cell-cell interactions, and nuclei are stained with Hoechst 33342. Scale bar: 25 µm.

### Morphological characterisation of fallopian tube cells in a microfluidic chip

Having established that fallopian tube epithelial cells form extensive monolayer sheets in the chip, we next characterised their morphology and benchmarked it against cells grown in standard tissue culture (TC) wells. Nuclei were segmented using CellProfiler, and nearest-neighbour distance, eccentricity and solidity were measured for chip-grown (n = 292) and well-grown (n = 641) nuclei (Figure 5).

**Fig. 5.**
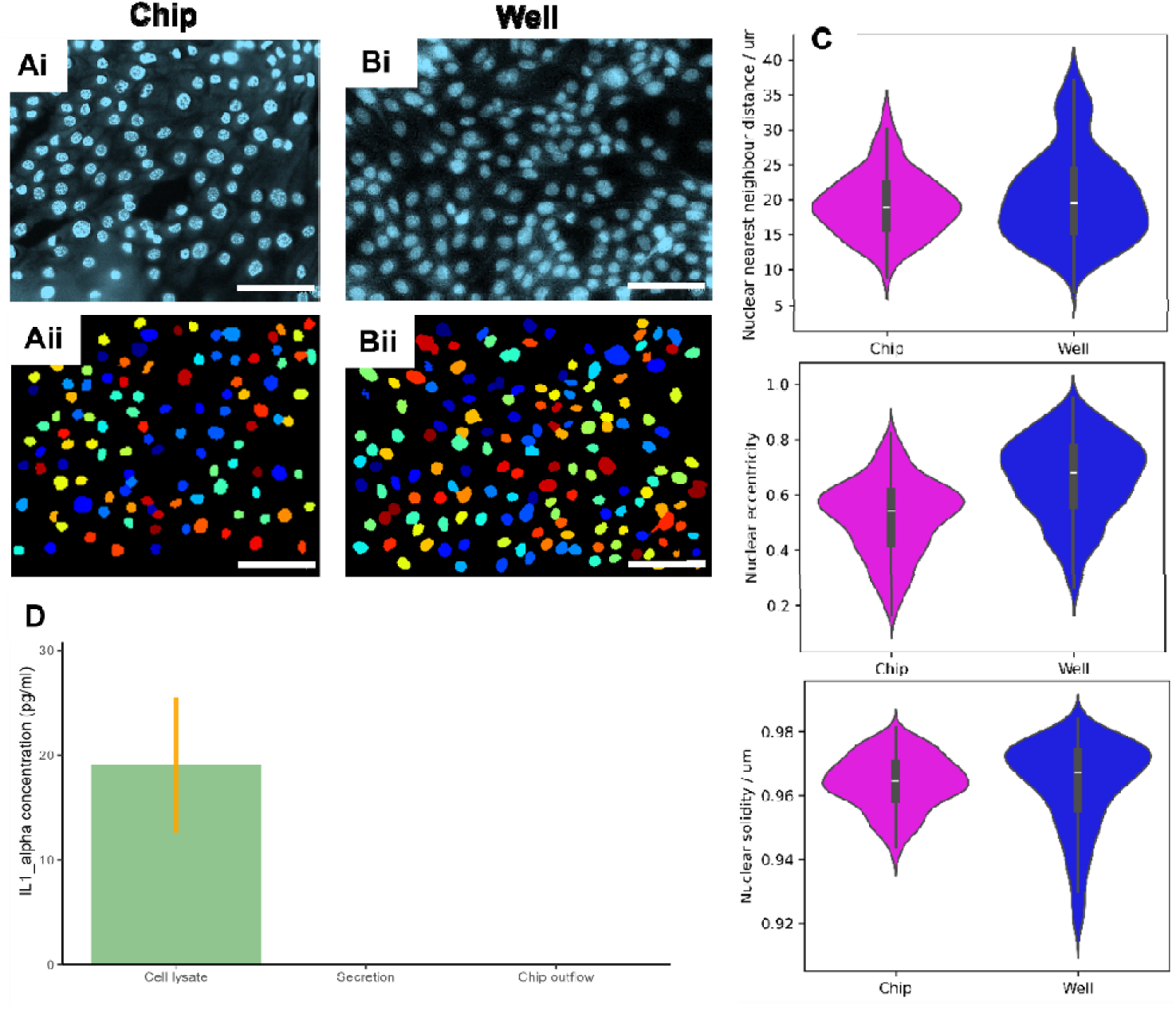
Morphology and inflammatory status of fallopian tube epithelium on-chip compared with standard culture. (Ai) Representative fluorescence image of Hoechst 33342-stained nuclei in fallopian tube epithelium grown on-chip. (Aii) Corresponding CellProfiler nuclear segmentation, with each segmented nucleus shown in a different colour. (Bi) Representative fluorescence image of Hoechst 33342-stained nuclei in fallopian tube epithelium grown in standard tissue-culture wells. (Bii) Corresponding CellProfiler nuclear segmentation, with each segmented nucleus shown in a different colour. (C) Violin plots of nuclear nearest-neighbour distance (Ci), eccentricity (Cii) and solidity (Ciii) for chip culture (magenta; n = 292 nuclei from three independent devices) and well culture (blue; n = 641 nuclei from three independent wells). Boxes show the interquartile range and white lines show the median. (D) IL-1α concentration measured by ELISA in fallopian tube organoid cell lysates, secretions from intact organoids, and day 7 outflow from opened organoids cultured on-chip. Cell lysates contained detectable IL-1α, whereas IL-1α in both intact-organoid secretions and chip outflow was below the assay detection limit. Scale bars: 50 µm.

Nearest-neighbour distance was similar in both formats, with a median of approximately 19 µm, but was more broadly distributed in well culture, indicating more variable cell packing. Eccentricity, a measure of elongation in which 0 represents a perfect circle and 1 a line, was lower on-chip (median approximately 0.54) than in wells (approximately 0.68), showing that chip-grown nuclei were rounder and less elongated. Solidity, the ratio of an object’s area to that of its convex hull, was close to 1 in both formats (median approximately 0.96–0.97), indicating regular nuclear outlines with few indentations. However, the well distribution had a longer tail towards lower values, reflecting a subpopulation of irregularly shaped nuclei that was largely absent on-chip. Together, these data indicate that the chip supports a more uniform epithelial layer, with more consistent nuclear shape and cell packing than standard well culture.

### Cell health on-chip assessed by IL-1α release

To assess whether mechanically opening organoids and culturing the resulting sheets under flow was associated with increased IL-1α release, we measured interleukin-1α (IL-1α), a damage-associated alarmin that is stored intracellularly and can be released when cells are stressed or lose membrane integrity. IL-1α was quantified by ELISA in three sample types: day 7 outflow from organoid-on-chip devices; secretions from intact fallopian tube organoids cultured in Matrigel domes, which served as a reference; and lysates of fallopian tube organoids, which represent the total intracellular IL-1α pool (Figure 5D). IL-1α was readily detected in organoid lysates (19.0 ± 6.4 pg/ml, mean ± SD, n = 7), confirming that the cells contain a measurable intracellular pool of IL-1α. In contrast, IL-1α was below the assay’s limit of detection (1.1 pg/ml) in both chip outflow and intact-organoid secretions. Thus, culturing mechanically opened organoids as sheets on-chip did not result in a detectable increase in IL-1α release relative to intact organoids. As both extracellular sample types were below the detection limit, these data do not establish equivalence between their secretion profiles.

### Modelling early tubal disease with tumour spheroid co-culture

To test whether the platform could support disease-relevant co-culture, GFP-tagged OVCAR3 ovarian cancer spheroids were seeded into a microfluidic device containing an established fallopian tube epithelial sheet. Immediately after inoculation (D0), spheroids appeared as discrete, compact clusters of GFP-positive cells on the epithelial sheet (Figure 6Ai). By 24 hours (D1), the GFP-positive signal was dispersed across a broader area, with fewer tightly compact clusters and more scattered individual cells (Figure 6Aii). This single-device proof-of-concept shows that the platform can support short-term co-culture of tumour and tubal epithelial compartments. Replicated experiments and quantitative measurements will be required to determine whether the observed redistribution represents attachment, migration, invasion, or another form of tumour–epithelium interaction.

**Fig. 6.**
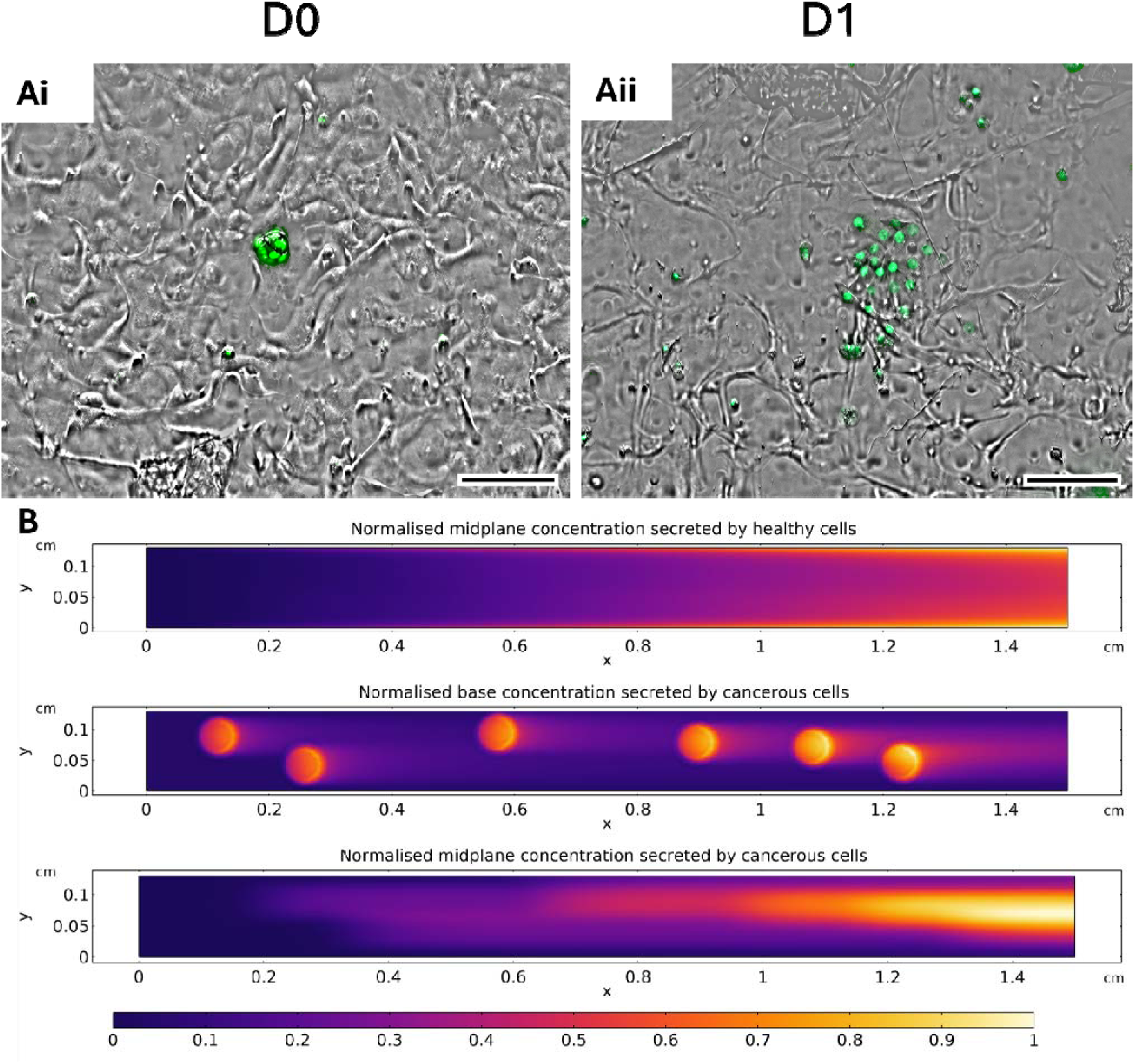
Proof-of-concept co-culture of ovarian cancer spheroids with a fallopian tube epithelial sheet and simulated distribution of secreted factors. OVCAR3 ovarian cancer spheroids were seeded into a microfluidic device containing a fallopian tube epithelial sheet. **(A)** Epifluorescence and brightfield images acquired immediately after spheroid inoculation (D0; Ai) and 24 hours later (D1; Aii). GFP-tagged OVCAR3 cells are shown by epifluorescence overlaid on the brightfield images. Images were collected from one chip using a 20× air objective; this experiment demonstrates technical feasibility and was not used for quantitative inference. Scale bars are 100 µm, except Ai, for which the scale bar is 50 µm. **(B)** Simulated transport under left-to-right flow along the 1.5 cm channel, shown as normalised concentration from 0 to 1. Top: concentration at the channel midplane for factors secreted by healthy fallopian tube cells. Middle: concentration at the channel base for factors secreted by six illustrative cancer-associated source regions, showing localised downstream plumes. Bottom: concentration at the channel midplane, where the plumes merge into a central band and accumulate downstream. The six simulated source regions illustrate spatial transport and are not a one-to-one reconstruction of the experimental spheroid distribution.

To predict how factors released by such lesions would be distributed within the device, we modelled secretion from healthy epithelium and from discrete tumour sites under flow (Figure 6B). Factors secreted uniformly by healthy epithelium accumulated progressively along the channel, reaching their highest concentration towards the outlet. Factors secreted by tumour sites formed localised plumes at the channel base that extended downstream of each site, and at the channel midplane these plumes merged into a central band that accumulated towards the outlet. Tumour-derived factors released anywhere along the channel are therefore delivered to the outlet, where they can be sampled in the effluent, while epithelium downstream of a lesion is exposed to its secreted factors. This supports use of the platform to study paracrine effects of early lesions on neighbouring tubal epithelium.

## Discussion

Our organoid-on-chip platform bridges static 3D organoid culture and dynamic microfluidic systems. Open loading ensures uniform cell seeding, giving more than 90% channel coverage by day 6 (n=5), and planar epithelial sheets facilitate high-resolution imaging that is difficult to achieve in an intact 3D organoid. Because the epithelium is derived from expandable organoid stock rather than freshly isolated tissue, the platform can support long-term, longitudinal experiments without repeated access to patient material, addressing a key practical barrier to fallopian tube organ-on-chip work. Morphological benchmarking of the organoid-derived epithelium against standard tissue culture (TC) wells suggests the chip supports a more uniform, cohesive epithelial layer than conventional culture. IL-1α was detectable in organoid lysates but below the assay limit of detection in both chip outflow and intact-organoid secretions. This indicates that mechanically opening the organoids and culturing them under flow did not produce a detectable increase in IL-1α release relative to intact organoids, although it does not demonstrate that the broader secretion profiles of the two culture formats are equivalent. This accessibility may also help address a recognised barrier to wider organ-on-chip adoption, namely that complex fabrication and operation requirements have tended to confine the technology to specialist engineering laboratories rather than mainstream biomedical and clinical settings [19].

Beyond its intended use for studying tubal damage from infection, endometriosis and occlusion, we show as a proof of concept that the platform can support co-culture with OVCAR3 ovarian cancer spheroids, a first step towards modelling early tumour-epithelium interactions relevant to gynaecological cancer. Flow modelling further indicates that factors secreted by discrete lesions are carried downstream and concentrated towards the outlet, so that lesion-derived signals can be sampled in the effluent while their effects on downstream epithelium are imaged. This is particularly relevant to the STIC-to-HGSOC pathway: STIC is a small, often microscopic lesion typically confined to the distal fimbrial epithelium and defined histologically by aberrant p53 expression alongside a high Ki-67 proliferative index, and is now widely accepted as the precursor from which most HGSOC arises [21]. Precisely because it is small and usually asymptomatic, STIC is detected almost exclusively as an incidental finding in prophylactic salpingectomy specimens, with a pooled incidence of around 7% even in this high-risk population [22]; prospective, longitudinal study of the tumour-epithelium interface as it forms is therefore essentially inaccessible in patients. The tractability and imageability demonstrated here indicate that the platform is not limited to this one application, and could similarly be extended to other early disease processes affecting the tube, from infection and inflammation-driven damage through to other adnexal malignancies, with potential future application to drug and therapeutic screening more broadly. A natural extension specific to STIC modelling would be to introduce defined TP53 mutations into the fallopian tube organoids themselves, rather than relying on an already-transformed tumour cell line, to more directly model de novo STIC lesion formation within the native epithelium.

As with any first-generation organ-on-chip platform, there is clear scope for further development. The device already incorporates a dedicated vasculature-mimicking compartment that runs parallel to the cell-populated epithelial channel and is perfused via manual pipette tips. This vascular compartment is not currently lined with cells, however, and its perfusion is not yet matched to the physiological flow parameters of the native fallopian tube vasculature. A natural next iteration would be to line this channel with endothelial cells and to calibrate its perfusion more closely to *in vivo* flow, requiring nutrients, metabolites and secreted factors to cross a functional endothelial barrier as they would through native tubal vasculature, rather than passing between compartments unmediated as at present, and further increasing the representative nature of the model. Immune cell components and cyclical hormone exposure also shape fallopian tube physiology *in vivo* and are not yet incorporated here, and functional readouts such as ciliary beat frequency were not directly quantified in this study. Other female reproductive tract chips indicate these are tractable additions rather than fundamental barriers: compartmentalised co-culture with endothelial cells has been used to model vascular-stromal interactions and cyclical hormone exposure in an endometrium-on-chip [20], suggesting a route by which comparable features could be incorporated into future iterations of this platform. Results to date are also based on organoids expanded from cryopreserved stock rather than on fresh tissue from multiple donors, and validating reproducibility across organoid lines from a larger, more diverse donor biobank will be an important next step before the platform is used for applications such as patient-specific disease modelling or drug screening.

## Accessibility and Versatility

Perfusion is driven passively by the difference in medium volume between filtered P200 pipette tip reservoirs at the inlet and outlet ports, so no pumps, tubing or flow controllers are required. This low-cost approach means the platform can be adopted by laboratories with limited engineering infrastructure. Pump-driven perfusion could be incorporated where more precise control of shear stress is needed, but was not used in this study.

## Methods

### Device Fabrication

PDMS microfluidic devices were fabricated by casting prepared PDMS into a negative silicone wafer master mould. Reactive plasma etching was used to define the mould channels. PDMS was prepared using Sylgard-184 silicone elastomer kit (Dow, Michigan, US) with a 10:1 ratio of base to curing agent. Once mixed the PDMS was degassed in a vacuum chamber for 45 minutes to remove air bubbles. PDMS was then poured into the master mould and cured at 80°C for 30 minutes in an oven. Once cured the PDMS was peeled away from the master mould and 1mm ports punched in the inlets and outlets. The side of the PDMS device containing the etched channels was then covered with Scotch tape and placed in a container until required. For those microfluidic devices bonded to glass coverslips, both the PDMS device and glass coverslips underwent plasma activation (O, 30 W, 60 s) in an HPT-100 plasma system (Henniker Plasma, Cheshire, UK). The underside of the PDMS device containing the channels was placed on the plasma treated glass coverslip and pressed gently to bond. To strengthen the bonding between PDMS and glass, the devices were placed in an oven at 80°C for 30 minutes. Bonded devices were covered until required.

### Cell and Organoid Preparation

Fallopian tube samples used to generate the organoids were provided by Richard Edmondson. Organoids were expanded from cryopreserved stock originally derived from fallopian tube tissue collected with written informed consent under [ethics approval or biobank reference]. The OVCAR3 cell line was provided by Christine Schmidt, who also advised on its use.

Fallopian tube organoids were cultured in serum free advanced DMEM/F12 (Thermo Fisher, Massachusetts, US) containing 1x penicillin-streptomycin-glutamine (PSG) according to the manufacturer’s instructions (Thermo Fisher), growth factors and other additives adapted from Kessler et al. [14]. Media was supplemented with 12 mM HEPES (Sigma-Aldrich, Missouri, US), 200 ng/ml recombinant Noggin (Thermo Fisher), 100 ng/ml recombinant human Fibroblast Growth Factor-10 (Thermo Fisher), 10 ng/ml recombinant Epidermal Growth Factor (PeproTech) 1mM Nicotinamide (Sigma-Aldrich) 100 ng/ml Wnt3A (Thermo Fisher), 10 µM ROCK inhibitor (Y-27632) (Sigma-Aldrich) and 0.5µM TGF-β R Kinase Inhibitor (SB431542) (Sigma-Aldrich). Additionally, 1% N2 supplement (Gibco), 2% B27 supplement minus vitamin A (Thermo Fisher) and 1.25mM N-Acetyl-L-cysteine (Sigma-Aldrich) were added to the media.

Fallopian tube organoids were grown in 24 well plates suspended in 50 µl of Matrigel (Corning, New York, US) diluted 1:5 in FT cell culture media on sterile glass coverslips. Organoids were passaged once a week to ensure larger organoid spheres were produced and split to a ratio of 1:2 or 1:1.5 dense wells to new wells for maintenance of organoids. Media was changed every 3-4 days. To passage organoids, media was removed from wells and the Matrigel broken mechanically by pipetting with a P1000 pipette and were transferred to microcentrifuge tubes and centrifuged at 2,400 x g for 6 minutes. Once pelleted, supernatant was removed and organoid pellets resuspended in basal media (advanced DMEM/F12 + PSG) and pipetted up and down 300 times to fragment. Resuspended cells were centrifuged at 600 x g for 6 minutes. Supernatant was again removed from cell pellets and cells resuspended in basal media and pipetted up and down 100 times to further fragment. Organoids were centrifuged once again at 1200 x g for 6 minutes. Finally, the supernatant was removed, and cell pellet resuspended in 60 µl of 1:5 Matrigel solution (Corning) and kept on ice. 50 µl of organoids resuspended in 1:5 Matrigel solution were then seeded into new 24 well plate wells with sterile glass coverslips and Matrigel allowed to solidify at 37°C for 30 minutes. Complete media was then pipetted into each well.

OVCAR3 cells were cultured in DMEM (Thermo Fisher) supplemented with 10% FBS and PSG at 37 °C and 5% CO in T75 tissue culture flasks (Corning). OVCAR3 cells were passaged every 3–4 days using TrypLE (Thermo Fisher). OVCAR3 spheroids were generated using an AggreWell 400 plate (STEMCELL Technologies). The AggreWell plate was treated with anti-adhesive solution (STEMCELL Technologies) and washed three times with PBS to remove excess solution. To generate OVCAR3 spheroids with a diameter of approximately 50 µm, 60,000 cells were seeded into each AggreWell 400 well with 2 mL of basal medium and incubated at 37 °C and 5% CO for 24 hours. OVCAR3 spheroids were harvested by aspirating the medium, gently pipetting up and down, and transferring the suspension to a microcentrifuge tube. Spheroids were pelleted by centrifugation at 900 × g for 6 minutes and gently resuspended in 100 µL of basal medium. To confirm that spheroids remained intact, 10 µL of the suspension was placed on a glass coverslip and examined by brightfield microscopy.

### Preparation of microfluidic devices

PDMS microfluidic devices and glass coverslips were first soaked in 70% ethanol to disinfect and allowed to air dry. The channels of the microfluidic devices and glass coverslips were coated with the cell and tissue adhesive CellTak (Corning, New York, US). CellTak was prepared according to the manufacturer’s instructions. To prepare 300 µl of neutralised CellTak, 260 µL of sterile 0.1 M NaHCO (Sigma-Aldrich) at pH 8.0, 30 µL of Cell-Tak stock (1 mg/mL in 5% acetic acid) and 10 µL of 1 N NaOH (Sigma-Aldrich) were mixed and used immediately (within 10 minutes) to avoid precipitation. CellTak was incubated at room temperature for 30 minutes and then removed, with washing of the device and glass coverslip with distilled water.

Microfluidic devices bonded to glass coverslips were similarly flushed with 70% ethanol and then rinsed with distilled water. CellTak (Corning) was flushed through the device and incubated at room temperature for 30 min. Distilled water was flushed through the device to rinse excess CellTak (Corning).

## Computational flow modelling

Due to the small scale of the device, inertia is negligible and the interior flow can be modelled by the incompressible Stokes equations. As the device height is small compared to the width and depth, we can use a depth-averaged approach as discussed by Ding et al. [23], where the flow is purely horizontal, and takes a parabolic flow profile across this thin direction but with a horizontal vector amplitude to be determined. There is no twisting of the flow field through the depth. The horizontal vector amplitude is zero on all no-slip boundaries of the device (i.e. side walls, pillars and posts), adjusting over a distance comparable to the device height according to 2D Brinkman-like PDE [23]. We solve this PDE using code from Harris [24], and compute 2D streamlines in Paraview.

Two cell-lined channel-base configurations were modelled. In the first, the entire base was lined with healthy cells with a spatially uniform secretion flux. In the second, six randomly distributed circular cancer-associated source regions collectively occupied 10% of the base area, with healthy cells occupying the remainder. For both configurations, the velocity field was calculated in COMSOL Multiphysics by solving the Navier–Stokes equations with no-slip conditions at the channel walls, a fully developed flow condition at the inlet and zero pressure at the outlet. The resulting velocity field was supplied to the Transport of Diluted Species module to solve the reaction–advection–diffusion equations for substances released at the channel base. In the mixed configuration, cancer-associated secretion fluxes were imposed only within the six circular source regions, while healthy-cell secretion fluxes were imposed over the remaining base. Concentrations were normalised to their respective maximum values. Concentration distributions were examined at the channel base and midplane.

## Preparation of cells for microfluidic inoculation

To prepare fallopian tube organoids for inoculation into microfluidic devices, organoids were broken mechanically following the passaging protocol above once organoids had been grown for one week. Organoid fragments were resuspended in 1:5 Matrigel solution. A ratio of 1.5 wells of organoids was inoculated per device at a volume of 10 µl per device.

OVCAR3 spheroids were generated using the above protocol and then injected into a microfluidic device containing the fallopian tube organoid monolayer at day 7 post fallopian tube inoculation. OVCAR3 spheroids were injected via the input port gently to maintain the spheroid structure once in the microfluidic device. 100 µl of resuspended OVCAR3 spheroids were injected.

## Open microfluidic inoculation with fallopian tube organoids

PDMS microfluidic devices were inoculated with fallopian tube organoids by pipetting fragmented organoids onto the underside of the PDMS microfluidic previously coated with CellTak. A total volume of 10 µl of fallopian tube organoid fragments suspended in 1:5 Matrigel solution (Corning) were spotted in 2 µl dots along the PDMS microfluidic coated with CellTak. PDMS microfluidic devices were placed carefully on top of the inoculated glass coverslip. The device was attached to the glass coverslip by sealing the PDMS edges with silicone gel and incubated at 37 °C for 30 minutes. Once sealed, P200 pipette tips containing 100 µl of advanced DMEM: F12 with 10 % foetal bovine serum were inserted into the outlet ports and P200 tips containing 200 µl of the same media inserted into the inlet ports and gently pushed using a pipette. This was to achieve slow passive flow from inlet to outlets ports through the device. Media was changed every 3-4 days by removing the inlet port pipette tips and replaced with fresh media. For longitudinal imaging of epithelial sheet formation and maintenance, cells were cultured in the device for up to 10 days post inoculation (Figure 3). Effluent was collected daily from each chip through day 10 by removing outlet port tips and then frozen at –20 °C.

## Imaging

Bright-field microscopy (20x/0.75 NA objective) was conducted daily on microfluidic devices inoculated with fallopian tube organoids, including immediately after inoculation (D0). The length of the microfluidic device was inspected for cell distribution and growth.

Once sufficient growth was achieved in the devices, fluorescence microscopy was performed (20x air, 40x/1.2 NA, 60x and 100x/1.4 NA objectives) after staining cells with SiR-Actin (Spirochrome, Switzerland; cat. no. SC001) and Hoechst 33342 (Thermo Fisher Scientific, USA; cat. no. H3570). Images were analysed using FIJI and CellProfiler.

### Sheet coverage

Coverage was quantified from brightfield images spanning the full channel length, acquired daily. Epithelium-covered regions were segmented in FIJI by thresholding and using a top hat mask, and coverage was expressed as the percentage of the cell culture channel area occupied by epithelium (n = 5 devices).

### ZO-1 immunostaining

Devices were fixed on-chip with 4% paraformaldehyde for 30 min at room temperature, permeabilised with 0.1% Triton X-100 for 30 min and blocked with 1% BSA in PBS for 2 h at room temperature, with all solutions introduced via the inlet ports and devices flushed with PBS between steps. Cells were incubated with anti-ZO-1 primary antibody (ZO-1 Polyclonal Antibody, Invitrogen, cat. no. 61-7300, 1:200) overnight at 4 °C, followed by Goat anti-Rabbit IgG (H+L) Secondary Antibody, Alexa Fluor 488, Invitrogen, cat. no. A-11008, 1:500 for 2 h at room temperature.

## Image analysis

### CellProfiler image processing and segmentation

Images were processed using FIJI. The brightness and contrast of the images was adjusted and some images were cropped to select regions with non-overlapping cells to allow segmentation. Nuclear segmentations were performed using the CellProfiler IdentifyPrimaryObjects module. Eccentricity and solidity were obtained using the MeasureObjectSizeShape module and nearest neighbour distances using the MeasureObjectNeighbors module.

### Data analysis

Violin plots were plotted from the segmentation data obtained from CellProfiler. Outliers were removed, calculated as data points below Q1 – 1.5 x IQR or above Q3 + 1.5 x IQR. For chip data n = 292 and for well data n = 641.

### Statistical analysis

Nuclear morphology data are shown as violin plots of all segmented nuclei (chip, n = 292 nuclei from 3 independent devices; well, n = 641 nuclei from 3 independent wells), with medians and interquartile ranges indicated. Sheet coverage is reported for n = 5 independent devices. IL-1α concentrations are presented as mean ± SD (n = 7 lysate samples). IL-1α in chip outflow and intact organoid secretions was below the assay limit of detection, so no statistical comparison was performed. Analyses were performed in Python (SciPy and seaborn).

### Biochemical Assays

To determine whether opening organoids and culturing them on-chip induces cellular damage or inflammation, interleukin-1α (IL-1α) was measured by ELISA. IL-1α is a damage-associated inflammatory marker held intracellularly and released on cell stress or necrosis. It was measured in three sample types: outflow from microfluidic devices containing opened fallopian tube organoid sheets, collected at day 7; secretions harvested from intact fallopian tube organoids; and cell lysates of fallopian tube organoids. The cell lysates represent the total intracellular IL-1α pool and served as a reference for released IL-1α. Samples were stored at −80 °C (lysates and organoid secretions) or −20 °C (chip outflow) until analysis and thawed once before use. IL-1α was quantified using the Human IL-1 alpha ELISA Kit (Invitrogen, Thermo Fisher Scientific; cat. no. BMS243-2) according to the manufacturer’s instructions, with samples assayed neat in triplicate. Absorbance was read at 450 nm, and concentrations were calculated from a standard curve fitted with a four-parameter logistic model. The assay range was 1.6 to 100 pg/ml with an analytical sensitivity of 1.1 pg/ml, and samples below this limit were reported as not detected.

Secretions were harvested from fallopian tube organoids by removing the culture medium, treating each well with cell-recovery solution, and incubating at 4 °C for 1.5 hours. The resulting organoid suspension was transferred to microcentrifuge tubes and centrifuged at 200 × g for 6 minutes. The supernatant was removed, and the organoids were washed with serum-free DMEM/F12 and centrifuged again at 200 × g for 6 minutes. After removal of the wash, 200 µL of serum-free medium was added. Samples were then centrifuged at 12,400 × g for 6 minutes, and the resulting supernatant was collected as the organoid secretion sample and stored at −80 °C. Each secretion sample represented material from two wells of organoids. The resulting organoid pellets were also stored at −80 °C for subsequent lysate preparation. For lysate preparation, pellets were thawed and incubated on ice for 15 minutes in 100 µL of radioimmunoprecipitation assay (RIPA) buffer containing protease and phosphatase inhibitors, with occasional vortex mixing.

## Competing interests

R.T. is the founder of InViFlo, a University of Manchester spin-out company developing microfluidic organ-on-chip platforms, and is an inventor on patent application PCT/GB2025/052644 relating to this technology. The remaining authors declare no competing interests.

## Funding declaration

This work was funded by the University of Manchester Dame Kathleen Ollerenshaw Fellowship and the University of Manchester Research Institute (UMRI).

## Acknowledgements

We thank Stephen Taylor (Manchester) for kindly providing the GFP-H2B OVCAR3 cell line[1]. We also thank Richard Edmondson for providing the fallopian tube samples used to generate the organoids and for his guidance on developing a productive fallopian tube model. C.K.S. acknowledges funding by the MRC (research grant MR/X008754/1) and CRUK (ACED-funded NEMO (Novel Early Markers of Ovarian cancer) programme grant ACEPGM-2023/100001).

